# Altered hierarchical gradients of intrinsic neural timescales in mild cognitive impairment and Alzheimer’s disease

**DOI:** 10.1101/2023.09.26.559549

**Authors:** Aiying Zhang, Kenneth Wengler, Xi Zhu, Guillermo Horga, Terry E. Goldberg, Seonjoo Lee, Alzheimer’s Disease Neuroimaging Initiative

## Abstract

Alzheimer’s disease (AD) is a devastating neurodegenerative disease that affects millions of older adults in the US and worldwide. Resting-state functional magnetic resonance imaging (rs-fMRI) has become a widely used neuroimaging tool to study neurophysiology in AD and its prodromal condition, mild cognitive impairment (MCI). The intrinsic neural timescale (INT), which can be estimated through the magnitude of the autocorrelation of intrinsic neural signals using rs-fMRI, is thought to quantify the duration that neural information is stored in a local cortical circuit. The heterogeneity of the timescales is considered to be a basis of the functional hierarchy in the brain. In addition, INT captures an aspect of circuit dynamics relevant to excitation/inhibition (E/I) balance, which is thought to be broadly relevant for cognitive functions. Here we examined its relevance to AD. We used rs-fMRI data of 904 individuals from the Alzheimer’s Disease Neuroimaging Initiative (ADNI) database. The subjects were divided into 4 groups based on their baseline and end-visit clinical status, which were cognitively normal (CN), stable MCI, Converter, and AD groups. Linear mixed effect model and pairwise comparison were implemented to investigate the large-scale hierarchical organization and local differences. We observed high similarities between AD and Converter groups. Specifically, among the eight identified ROIs with distinct INT alterations in AD, three ROIs (inferior temporal, caudate, pallidum areas) exhibit stable and significant alteration in AD converter. In addition, distinct INT related pathological changes in stable MCI and AD/Converter were found. For AD and Converter groups, neural information is stored for a longer time in lower hierarchical order areas, while higher levels of hierarchy seem to be preferentially impaired in stable MCI leading to a less pronounced hierarchical gradient effect. These results inform that the INT holds great potential as an additional measure for AD prediction, a stable biomarker for clinical diagnosis and an important therapeutic target in AD.

## 1 Introduction

Alzheimer’s disease (AD), as the most common form of dementia, is estimated to affect 6.5 million older adults in the Unites States, a number which could increase to 13.8 million by 2060 barring the development of medical breakthroughs to prevent, slow or cure AD (1). AD is a devastating neurodegenerative disease, beginning with mild memory loss and progressively leading to compromises in other cognitive domains and impairments in everyday function (2).

Resting-state functional magnetic resonance imaging (rs-fMRI) has become a widely used neuroimaging tool to study neurophysiology in AD and its prodromal condition, mild cognitive impairment (MCI), because of MRI’s relative simplicity of use, noninvasive nature, and relatively high spatial resolution. The rs-fMRI functional connectivity (FC) of brain networks refers to the inter-regional synchrony detected from the blood oxygen level dependent (BOLD) fMRI sequence. It is an emerging AD biomarker that holds promise for early diagnosis (3; 4). Multiple studies have reported reduced connectivity in the default mode network (DMN) (5; 6)– a network that includes lateral and medial prefrontal, lateral temporal lobe, superior parietal lobe that is considered a predictor of cognitive decline (7). The salience network– a network involving insula and anterior cingulate regions, is thought to be involved in the selection of goal directed behavior in patients with AD and MCI (8). However, FC does not reflect the fundamental organizational principles that characterize the functional architecture of the human brain (9; 10).

In contrast, the intrinsic neural timescale (INT), is thought to quantify the duration that neural information is stored in a local cortical circuit – a property that directly supports functional specialization (11). Such neural timescales demonstrate a distinct gradient in the brain (12), where densely interconnected associative regions, such as prefrontal and parietal areas, have longer timescales compared to peripheral sensory areas. This heterogeneity of timescales is considered a basis of the functional hierarchy in the brain. The INT can be estimated from the magnitude of the temporal autocorrelation in rs-fMRI signals (11; 13; 14). It has been shown that metrics using the temporal autocorrelation of rs-fMRI are reliable, causally linked to biological processes beyond a FC network framework, and relevant to neuropsychiatric disease including dementias (15).

INT captures an aspect of circuit dynamics relevant to excitation/inhibition (E/I) balance, which is thought to be broadly relevant for cognitive functions (16). Specifically, large-scale biophysical modeling has demonstrated that INT is dependent upon the strength of recurrent excitation —a biophysical property shown to support working memory and perceptual integration (17). Furthermore, several disorders with cognitive dysfunction (e.g., schizophrenia (14; 18), Parkinson’s disease (19) and epilepsy (20)) have shown alterations in INT and this aspect of circuit dynamics may also be relevant to other neuropsychiatric disorders such as AD. One of the pathological hallmarks of AD is the accumulation of amyloid-*β* (A*β*) peptides in the brain that occurs long before clinical disease onset, synaptic failure due to axonal terminal dysfunction (reduced action potentials) when adjacent to amyloid neuritic plaques alters the E/I balance (21). On the other hand, studies in mouse models suggest that network hyperexcitability can occur in the early stages of Alzheimer’s disease and contribute to cognitive decline (22). Given that, the pathological distribution of *Aβ* plaques and tau paired helical filaments generally spares primary somatosensory and motor cortices and primary visual cortex until very late in the disease process while compromising multiple association cortices, the normal hierarchical gradient of INT could potentially be altered (23; 24).

To examine alterations of E/I balance and the associated hierarchical gradient, we applied rs-fMRI to test whether distinct changes of INT at low and high levels of the neural hierarchies are present in people with MCI, people that will later progress to AD and AD patients, respectively.

## 2 Methods and Materials

### 2.1 Participants

Data used in this study were obtained from the Alzheimer’s Disease Neuroimaging Initiative (ADNI) database (adni.loni.usc.edu). ADNI was launched in 2003 as a public-private partnership. The primary goal of ADNI has been to test whether serial magnetic resonance imaging (MRI), positron emission tomography (PET), other biological markers, and clinical and neuropsychological assessments can be combined to measure the progression of mild cognitive impairment (MCI) and early Alzheimer’s disease (AD).

A detailed description of the ADNI cohort has been previously published (25). ADNI has recruited individuals that are cognitively normal (CN), have mild cognitive impairment (MCI), or have dementia or Alzheimer’s diseases (AD). We used the rs-fMRI data of 945 subjects at baseline. We excluded participants with poor data quality (see below for details) and the final sample included 904 participants. Further, we divided the subjects into 4 groups based on their baseline and end-visit clinical status (up to 10 years before they drop off the study): 1) CN group if the subject was CN at baseline and not demented at end visit; 2) MCI group if the subject was MCI at baseline and not demented at end visit; 3) Converter group if the subject was CN/MCI at baseline and demented at end visit; 4) AD group if the subject was demented at baseline. Demographic characteristics are shown in Table 1. Of the CN/MCI individuals who progressed to AD (called Converters hereafter), four were cognitively normal at baseline while the remainder were diagnosed as MCI at baseline.

**Table 1:** Age and Sex distribution by group.

|  | **CN**<br>(N=476) | **MCI**<br>(N=262) | **Converter**<br>(N=57) | **AD**<br>(N=109) | **Total**<br>(N=904) | p value |
| --- | --- | --- | --- | --- | --- | --- |
| **Age** |  |  |  |  |  | 0.006 |
| Mean | 73.2 | 73.5 | 75 | 75.9 | 73.7 |  |
| Min | 55.7 | 55.6 | 57.3 | 55.3 | 55.3 |  |
| Q1,Q3 | 67.7, 77.9 | 68.7, 79.1 | 70.2, 79.3 | 71.3, 81.8 | 68.4, 79.0 |  |
| Max | 95.5 | 93.3 | 89.7 | 96 | 96 |  |
| **Sex** |  |  |  |  |  | 0.002 |
| F | 281 (59.0%) | 122 (46.6%) | 24 (42.1%) | 54 (49.5%) | 481 (53.2%) |  |
| M | 195 (41.0%) | 140 (53.4%) | 33 (57.9%) | 55 (50.5%) | 423 (46.8%) |  |

### 2.2 Image acquisition and pre-processing

All MRI data were downloaded on August 18, 2020. T1-weighted (T1w) MRI and rs-fMRI data from 3T MRI scanners were included in this study.

T1w images were acquired with a gradient recalled echo (GRE) pulse sequence with acquisition in the sagittal plane with the following acquisition parameters: repetition time (TR) = 6.98 ms, echo time (TE) = 2.85 ms, inversion time (TI) = 400 ms, field of view = 26 cm, voxel size 1 *×* 1 *×* 1.2 mm and acquisition matrix = 256 *×* 256 *×* 196. All T1w images underwent automated quality control through MRIQC (26). For subjects with multiple available T1w images at the same visit, we selected the images with the best quality for further analysis (i.e., only one T1w image was selected for each subject). For all selected T1w images that passed the quality check, cross-sectional image processing was performed using FreeSurfer Version 7.1.1 (https://surfer.nmr.mgh.harvard.edu/). Region of interest (ROI)-specific cortical thickness (CT) values were extracted from the automated anatomical parcellation using the Desikan-Killiany Atlas (27). These CT values were used as covariates to examine the effect of atrophy, since cortical thinning has always been observed in neurodegenerative disorder (28) and it might potentially affect the INT values.

Rs-fMRI data were acquired with an echo-planar imaging sequence with the following acquisition parameters: 140 functional volumes, TR = 3000 ms, TE = 30 ms, flip angle = 80, number of slices = 48, slice thickness = 3.3 mm, voxel size = 3 *×* 3 *×* 3.3 mm and in plane matrix = 64 *×* 64. For each subject, the first 5 volumes of the functional images were discarded for signal equilibrium and to allow the participant’s adaptation to the scanning circumstances, leaving 135 resting-state volumes for further preprocessing. The preprocessing steps for rs-fMRI are described as follows. First, we used a custom methodology of fMRIPrep to generate a reference volume and its skull-stripped version. Head-motion parameters with respect to the BOLD reference including transformation matrices, and six corresponding rotation and translation parameters are estimated before any spatiotemporal filtering using mcflirt under FSL 5.0.9 (29). We then used 3dTshift from AFNI 20160207 for slice-time correction. The BOLD reference was co-registered to the T1w reference using bbregister by FreeSurfer, which implemented boundary-based registration with six degrees of freedom (30). Co-registration was configured with six degrees of freedom. We resampled the BOLD time-series into standard MNI152NLin2009cAsym space. Several confounding time-series metrics were calculated based on the preprocessed BOLD, i.e., the framewise displacement (FD) (31) and the root-mean-square difference (RMSD) of the time-series in the consecutive volumes (29). Contaminated volumes were then detected and classified as outliers by the criteria FD *>* 0.5mm or RMSD *>* 0.3% and replaced with new volumes generated by linear interpolation of adjacent volumes using the CONN toolbox. The three global signals are extracted within the cerebrospinal fluid (CSF), the white matter (WM), and the whole-brain masks. We further bandpass filtered the time-series with cut-off frequencies of 0.01 and 0.09 Hz. Finally, the covariates corresponding to head motion (6 realignment parameters), outliers, and the BOLD time series from the subject-specific WM and CSF masks were removed from the BOLD functional time series using linear regression. After estimating voxel-wise INT values (see Section 2.3) from the time-series, ROI-specific INT were calculated as the mean of the INT values of the voxels within the ROI from the automated anatomical parcellation using the Desikan-Killiany Atlas (27) for cortical areas, which results in 68 cortical ROIs (34 on each hemisphere), and the Aseg Atlas (32) for subcortical areas. The Desikan-Killiany Atlas has been widely used in previous AD studies since other refined brain parcellations does not fit well with aging brain, especially those with neurodegenerative disorder (28; 33), and it facilitates comparison to (and potential confound of) cortical thickness. We selected 16 subcortical ROIs (8 on each hemisphere), which includes thalamus, caudate, putamen, pallidum, accumbens area, hippocampus, amygdala and ventral diencephalon.

We excluded the participants 1) whose rs-fMRI contained *>* 30% frames flagged as motion outlier (27 subjects were excluded), 2) without diagnosis information (12 were excluded), or 3) had corrupted MRI images (2 were excluded). Therefore, 904 participants (age range: 55-96) were left for the analysis.

### 2.3 Intrinsic Neural Timescale (INT) Calculation

Before estimating the voxel-wise INT values, preprocessed rs-fMRI data were further processed with the following steps: (1) motion censoring of outliers that are consecutive in three and above volumes to be excluded and those volumes will be treated as NA; and 2) spatial smoothing with a 6 mm full-width-at-half-maximum Gaussian kernel. For each subject *i*, voxel *v*, the autocorrelation function (ACF) was estimated by

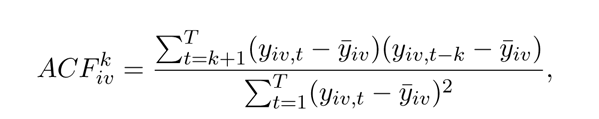

where *k* is the time lag, *k* = 1, 2*, . . ., T −* 1, *T* is the total number of timepoints, and **y***_iv_* = (*y*^1^ *, y*^2^ *, . . ., y^T^_iv_*) is the rs-fMRI signal sequence of voxel *v* for subject *i*. The voxel-wise INT was then estimated as the area under the curve of the ACF during the initial positive period:

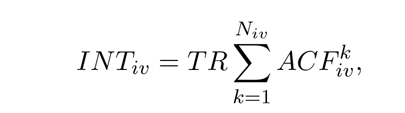

where TR is the repetition time, *N_iv_* is the lag directly proceeding the first negative ACF value of voxel *v* for subject *i*. Finally, the ROI-specific INT *INT_ij_* is calculated as the mean of the *INT_iv_*’s within ROI *j*.

### 2.4 Scanner Harmonization to Remove Site Effect

The ADNI data were collected from 58 sites. To remove undesired artifacts from scanner and site differences, we applied the ComBat harmonization method (34) to the INT and CT values. For each subject *i*, ROI *j*, and site *k* we fitted the following model:

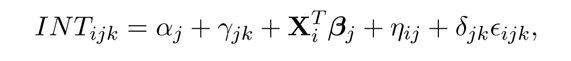

where **X***^T^* contains the covariates of interest, which includes age, sex, mean framewise displacement (MFD), and diagnosis group. The ComBat harmonized INT was

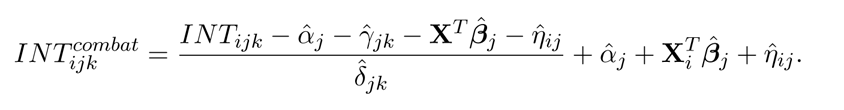

Similarly, we obtained the ComBat harmonized CT *CT ^combat^*.

### 2.5 Linear Mixed Effect Models to Detect Altered Hierarchical Gradient Effects on INT

Our previous study has validated INT as an index of cortical hierarchy using the Glasser MMP1.0 atlas (14). Here, to facilitate comparison with previous literature and with cortical thickness alterations, we defined the hierarchical level of the 68 ROIs by Desikan-Killiany atlas using the rs-fMRI of 100 unrelated young and healthy subjects from the Human Connectome Project (HCP) WU-Minn Consortium (35). Subcortical ROIs were excluded from hierarchical analyses given the disparate factors that shape the cortical hierarchy compared to subcortical hierarchies (where multiple hierarchies may exist within individual subcortical regions (36)). We used the average INT map in 2mm-isotroptic MNI space from 100 unrelated HCP young-adult subjects as in (14). The ROI-specific INT is calculated as the mean INT within each ROI. The hierarchical level is determined by ranking each ROI’s INT value from low to high corresponding to short to long INT, and unified to [0, 1] by dividing the total number of ROIs. We compared the hierarchical levels between the Glasser and Desikan-Killiany atlases as shown in Appendix B Figure A2, and confirmed that the Desikan-Killiany atlas maintains the hierarchical levels as the Glasser atlas. The exact hierarchical orderings of Desikan-Killiany atlas are visualized in Figure 1 a) and the full table is given in the Appendix C Table A3.

**Figure 1:**
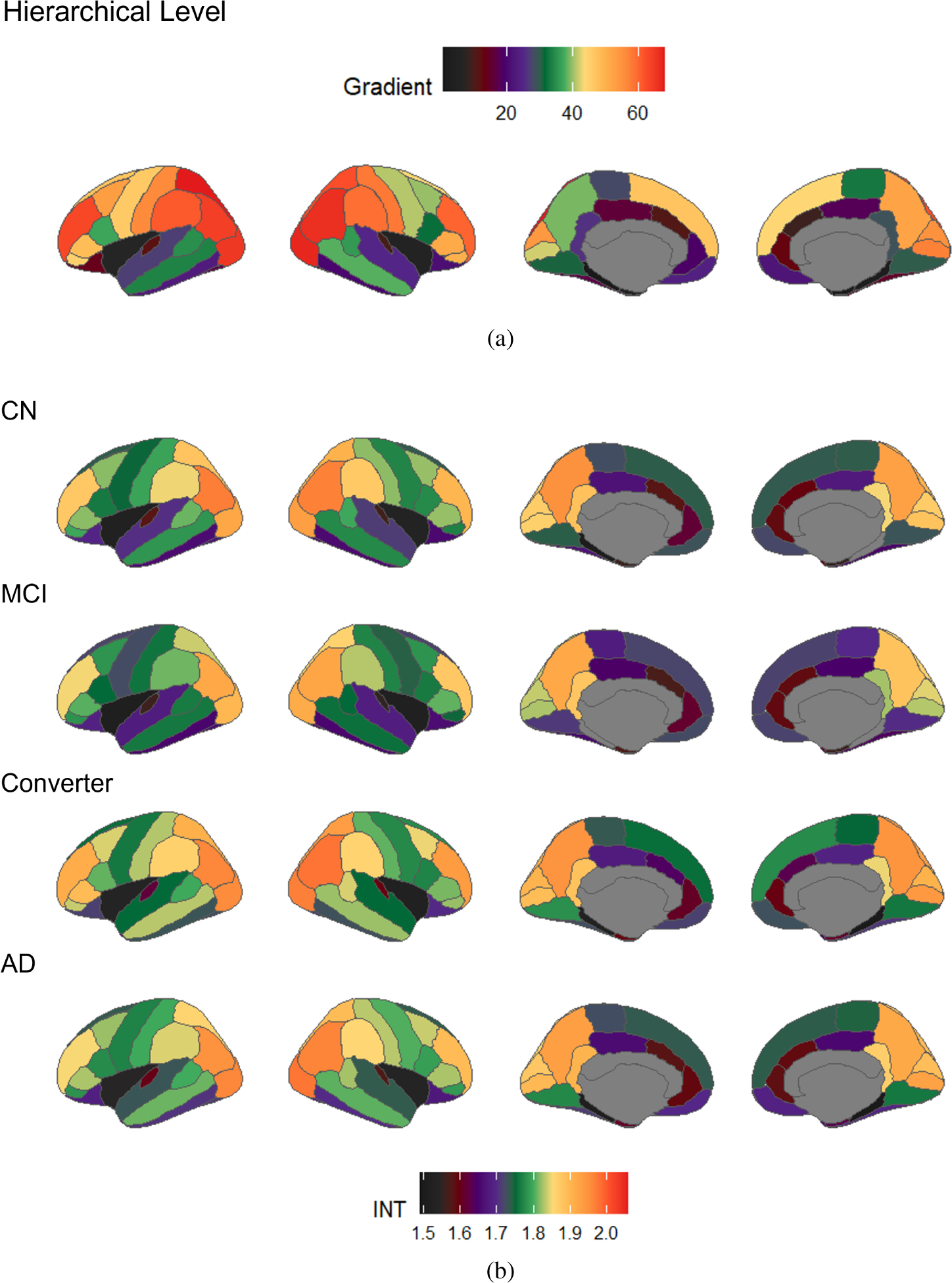
INT as a measure of brain hierarchical level. (a). The hierarchical gradients of the DKT atlas from the Human Connectome Project (HCP) dataset. (b). Parcellated group-averaged INT maps by groups from the Alzheimer’s Disease Neuroimaging Initiative (ADNI) database. The flame scale is ordered so that long (high) INTs are toward red and short (low) INTs are toward black.

We applied the linear mixed effect (LME) model to predict the INT value in our ADNI sample using hierarchical level (HL) in the cerebral cortex, assuming different intercepts and slopes by diagnosis group. We also considered age, sex, MFD, and CT as covariates (fixed effects), and allowed for variations of intercept and slope at the subject level (random effects). As such, for each subject *i*, and ROI *j*:

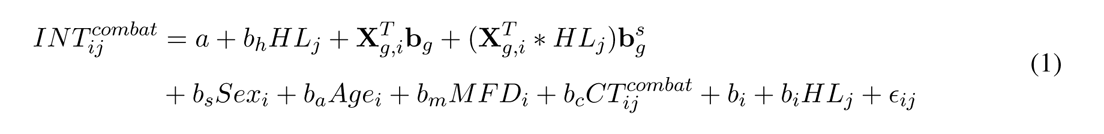

where *Sex_i_ = male)*, *X^T^_g,i_* = (*I(group_i_ = MCI), I(group_i_ = AD), I(group_i_* = *Progressor*)) is a vector that contains the diagnosis group information for subject *i*. T-statistics and the corresponding p-value were calculated for the regression coefficients, and their effect size (ES) was measured by partial *η*^2^, denoted as *η*^2^ (37).

### 2.6 Pairwise Comparison to Detect Significant INT Differences among Groups

We conducted pairwise comparison between diagnosis groups to identify significant INT differences. We averaged the INT values of the same ROIs on the left and right hemisphere, which left 34 cortical ROIs and 8 subcortical ROIs for comparison. The hypothesis testing was implemented through the following steps: first, ANOVA analysis was applied to screen for ROIs that have group differences using F-tests at a significance level *α* = 0.05; we included age, gender, MFD and ROI cortical thickness as covariates. Then, for the selected ROIs, t-tests were conducted to compare the differences between each pair of the 4 groups, i.e., CN, MCI, MCI converter and AD. Multiple comparison correction was performed using the Tukey method for comparing a family of 4 estimates, but not controlled for the 34 ROI comparisons. The effect size of the difference was measured by Cohen’s d.

### 2.7 Sensitivity Analysis

To test the robustness of the INT by hierarchical level, we did a series of sensitivity analysis and compared the results using equation (1) in Section 2.5. Specifically, we repeat the LME model 1) without removing site effects using ComBat for INT and cortical thickness, considering site is a potential source of artifacts in the neuroimaging data; 2) removing age from the covariates, since age and diagnosis are highly correlated which may attenuate the group effect; 3) removing cortical thickness from the covariates to examine the effect of atrophy; 4) including APOE4 — a strong risk factor gene for AD — as a covariate; 5) replacing the 4 groups with 3 groups (CN, MCI, and AD) only based on their baseline diagnosis; 6) excluding the 4 low ranked ROIs (entorhinal and parahippocampus) to avoid low hierarchical level areas implicated in AD pathology over-weighing the differences. Additionally, head motion is a common source of artifacts in fMRI data (31), we included MFD as a covariate in all analyses and investigate its effect on the results.

In the current dataset, 114 participants (12.6%) did not have APOE4 information, thus a subset of n=790 subjects were included for this analysis. Details can be found in Appendix A Table A1. The demographic information of the groups based on baseline diagnosis can be found in Appendix A Table A2.

### 2.8 Code and Data Availability

R code is available for INT calculation at the github website (https://github.com/Aiying0512/INT). Specifi-cally, we used the *acf* function which allows for NA/missing values. ComBat harmonization was implemented in R using the R package *neuroCombat*. Linear mixed effect models were implemented in R using the R package *lmerTest*. The pairwise comparison of multiple groups was implemented in R using the R package *emmeans*.

ADNI datasets are available to the research community upon request at www.adni.loni.usc.edu. The processed imaging data are available for the qualified investigators upon request at.

## 3 Results

### 3.1 Hierarchical Gradient Effect on Cortical INTs among Various Groups

Figure 1 shows the parcellated INT maps on the cerebral cortex for each of the 4 diagnostic groups and the 100 unrelated HCP subjects used to determine the cortical hierarchical levels along with the corresponding map of cortical hierarchical level. Following previous work, our main analysis focused on hierarchical gradient effects on INT (i.e., the relationship between INT values and hierarchical level).

The results of the estimated parameters in the LME model are shown in Table 2. The fitted lines of INT values as a function of hierarchical levels of the 4 diagnosis groups are visualized in Figure 2 a. This analysis only included the cortex given prior work demonstrating a separate hierarchical system in subcortical regions (36).

**Figure 2:**
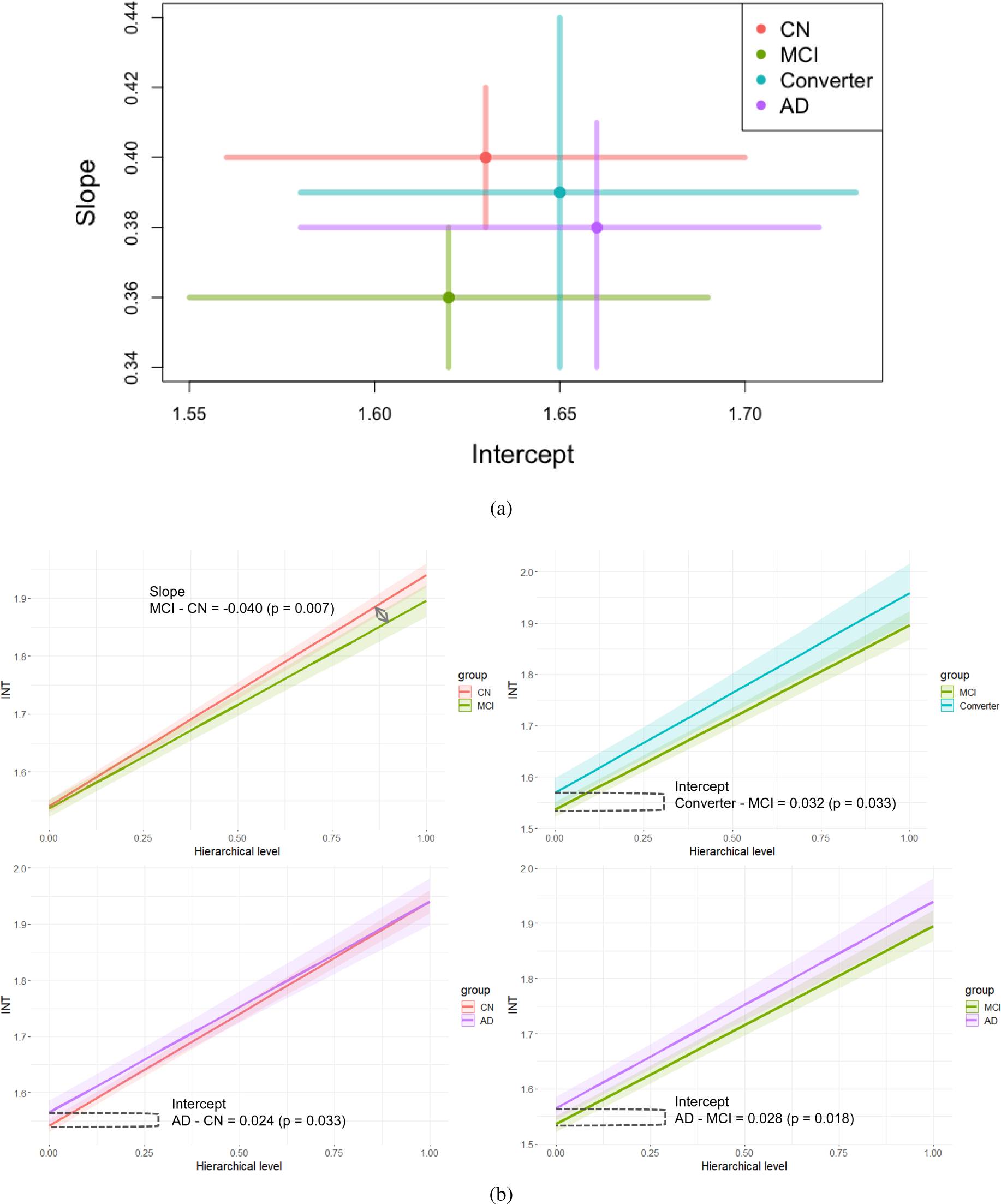
The results of the LME fitting. a) The estimated intercept and slope of the hierarchical gradient effects by diagnosis groups and the corresponding 95% confidence intervals. b) Pairwise comparisons between the groups with significant differences. INT values were plotted as a function of hierarchical level, as well as their 95% confidence intervals.

**Table 2:** Summary of the estimated parameters in the linear mixed effect model. CI: confidence interval; ES: effect size, which is measured by *η*^2^; df: degree of freedom.

| *Predictors* | *Estimates* | *95% CI* | **INT** |  |  |
| --- | --- | --- | --- | --- | --- |
| | | | t (df) | ES ($\eta_p^2$) | *p* |
| Intercept | 1.632 | 1.564 – 1.701 | 46.811 (984) | 0.690 | <**0.001** |
| Age at scan | -0.001 | -0.002 – -0.000 | -2.337 (898) | 0.006 | **0.019** |
| Sex [M] | 0.027 | 0.013 – 0.042 | 3.839 (897) | 0.016 | <**0.001** |
| Group [MCI] | -0.004 | -0.020 – 0.011 | -0.538 (897) | 0.000 | 0.59 |
| Group [Converter] | 0.028 | -0.001 – 0.057 | 1.920 (897) | 0.004 | 0.055 |
| Group [AD] | 0.024 | 0.002 – 0.046 | 2.133 (899) | 0.005 | **0.033** |
| HL | 0.400 | 0.381 – 0.416 | 44.450 (963) | 0.672 | <**0.001** |
| MFD | -0.012 | -0.103 – 0.079 | -0.270 (897) | 0.000 | 0.787 |
| Cortical Thickness | -0.004 | -0.008 – 0.001 | -1.649 (60472) | 0.000 | 0.099 |
| Group [MCI] * HL | -0.040 | -0.068 – -0.011 | -2.687 (900) | 0.008 | **0.007** |
| Group [Converter] * HL | -0.009 | -0.062 – 0.043 | -0.346 (900) | 0.000 | 0.729 |
| Group [AD] * HL | -0.024 | -0.064 – 0.016 | -1.166 (901) | 0.002 | 0.244 |

In terms of differences between diagnostic groups in their relationship between INT values and hierarchical level (see Figure 2 b and Table 3), the MCI group had a less pronounced hierarchical-gradient effect (as reflected in the contrasts of slope) when compared to the CN group in which INT values were similar in lower order areas but were shorter in higher order areas; the AD group showed longer INT values in lower order areas than CN group as reflected by the contrasts of intercept; the AD and Converter groups had longer INT across all cortical areas compared to the MCI group.

**Table 3:** Pairwise comparison of hierarchical gradient effects among diagnosis groups. ES: effect size, which is measured by Cohen’s d statistics; df: degree of freedom.

| *Contrasts* | **Intercept** |  |  |  | **Slope** |  |  |  |
| --- | --- | --- | --- | --- | --- | --- | --- | --- |
|  | Estimate | ES (d) | t(df) | *p* | Estimate | ES (d) | t (df) | *p* |
| CN - MCI | 0.004 | 0.036 | 0.538 (897) | 0.590 | 0.040 | 0.180 | 2.697 (900) | **0.007** |
| CN - Converter | -0.028 | -0.128 | -1.920 (897) | 0.055 | 0.009 | 0.023 | 0.346 (900) | 0.729 |
| CN - AD | -0.024 | -0.142 | -2.133 (899) | **0.033** | 0.024 | 0.078 | 1.166 (901) | 0.244 |
| MCI - Converter | -0.033 | -0.123 | -2.134 (897) | **0.033** | -0.030 | -0.072 | -1.088 (900) | 0.277 |
| MCI - AD | -0.028 | -0.158 | -2.361 (898) | **0.018** | -0.016 | -0.049 | -0.734 (900) | 0.463 |
| Converter - AD | 0.004 | 0.017 | 0.254 (897) | 0.800 | 0.014 | 0.031 | 0.460 (900) | 0.645 |

### 3.2 Assessment of Robustness of the INT by Hierarchical Gradient Effect

We set out to determine the robustness of differences between diagnostic groups in their relationship between INT values and hierarchical level through a series of sensitivity analyses. The p-values for linear mixed models are based on degree of freedom with Satterthwaite approximation. As shown in Figure A3 and Table A4 in Appendix D, the basic findings of INT by hierarchical level under various settings of sensitivity analyses are consistent with the main results in Figure 2, showing robustness to ComBat harmonization, alternative diagnostic group definitions, not covarying for age and cortical thickness, covarying for APOE4 status, and exclusion of the 4 lowest hierarchical-level cortical areas.

No significant effects of head motion (quantified as MFD) were observed in the main results (see Table A4) or any of the sensitivity analyses mentioned above (all p-values > 0.4). A significant age effect was observed in the main results (p-value = 0.019): INT was shorter as age increased when controlling for other covariates. When we excluded age to fit the LME model, the positive effect of AD group on INT was no longer significant (p-value =0.055 in Table A6), likely due to the close relationship between AD and age. We compared the main results and the ones without applying ComBat to remove site effects (see Appendix Figure A3,a and Table A4) and found site effects reduced the significance of age influence, suggesting that the removal of site effects reduced nonbiological variability in the neuroimaging data while maintaining meaningful biological variability. No significant cortical thickness effect was observed for hierarchical gradients of INT (although see below for effects in individual ROIs).

### 3.3 Cortical Thickness Effect on INT by ROIs

Although cortical thickness had no significant effect on the relationship between INT and hierarchical level, we were interested in investigating its effect on each individual ROI (averaged across hemisphere and adjusted for age, sex, MFD and diagnosis groups).

As shown in Figure 3, the cortical thickness of entorhinal (estimate = 0.046, *η*^2^ = 0.008, *t*(890) = 2.686, *p* = 0.007), insula (estimate= 0.1349, *η*^2^ = 0.010, *t*(896) = 2.932, p=0.028) and lingual (estimate= 0.083, *η*^2^ = 0.005, *t*(896) = 2.199, *p* = 0.003) areas had a positive relationship with INT, while the rostral middle frontal area (estimate= *−*0.1407, *η*^2^ = 0.005, *t*(896) = *−*2.069, *p* = 0.038) had a negative relationship with INT.

**Figure 3:**
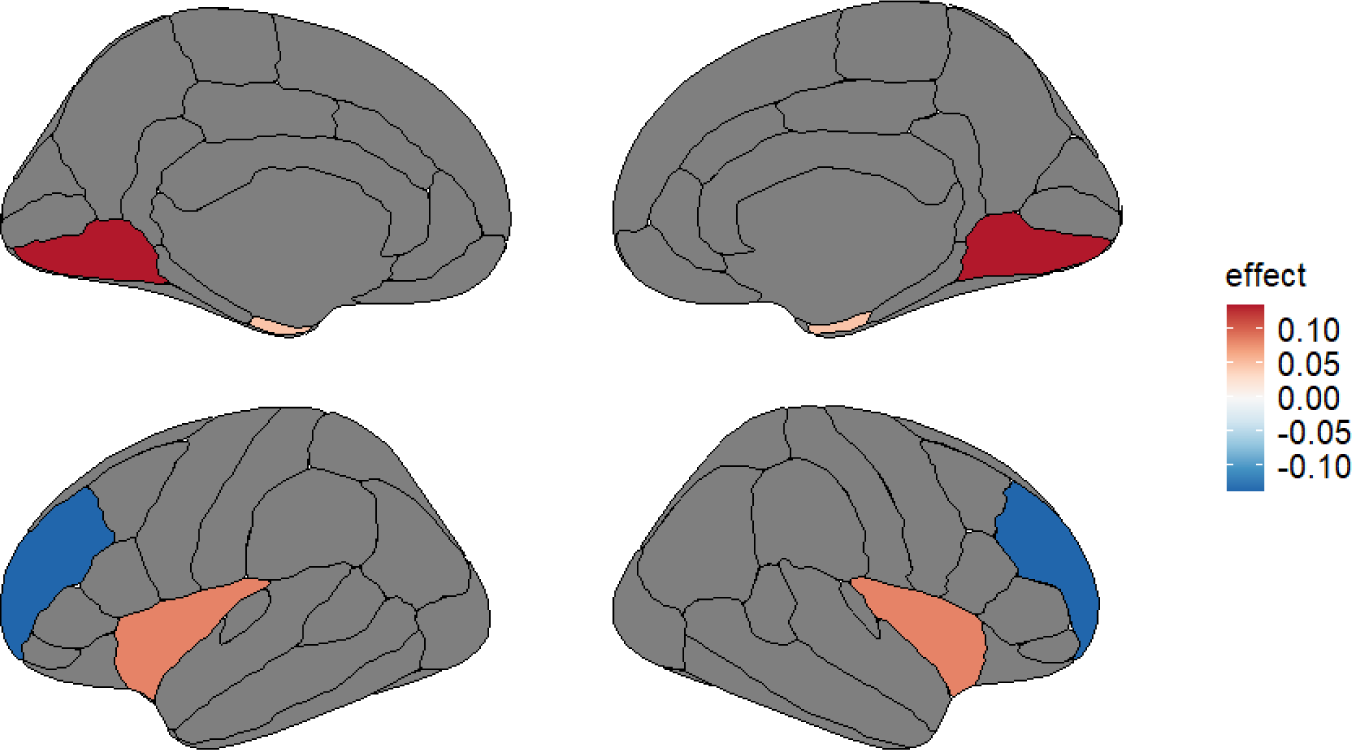
The identified ROIs that have significant cortical thickness effect on INT. The ROIs include: entorhinal, insula, lingual and rostral middle frontal areas. The colormap indicates the estimated regression coefficients.

### 3.4 Significant cortical ROIs in which AD and Converter groups had longer INT values than CN and MCI groups

Even though excluding low ranked ROIs (entorhinal cortex and parahippocampus) slightly attenuated the differences between diagnostic groups in their relationship between INT values and hierarchical level (Figure A3 (f)), the basic findings were still present. Therefore, it is critical to understand what ROIs drive the differences in these relationships. To this end, we conducted pairwise comparisons to examine the 34 cortical ROIs (averaged across hemisphere).

As shown in Figure 4, four ROIs were identified as having higher INT values in the AD group than the CN group: entorhinal (CN-AD: estimate= *−*0.086, Cohen’s *d* = *−*0.294, *t*(896) = *−*4.401, *p* = 0.0001), fusiform (CN-AD: estimate= *−*0.044, Cohen’s *d* = *−*0.174, *t*(896) = *−*2.612, *p* = 0.0452), inferior temporal (CN-AD: estimate= *−*0.086, Cohen’s *d* = *−*0.294, *t*(896) = *−*4.401, *p* = 0.0111) and temporal pole areas (CN-AD: estimate= *−*0.064, Cohen’s *d* = *−*0.210, *t*(896) = *−*3.143, *p* = 0.0094).

**Figure 4:**
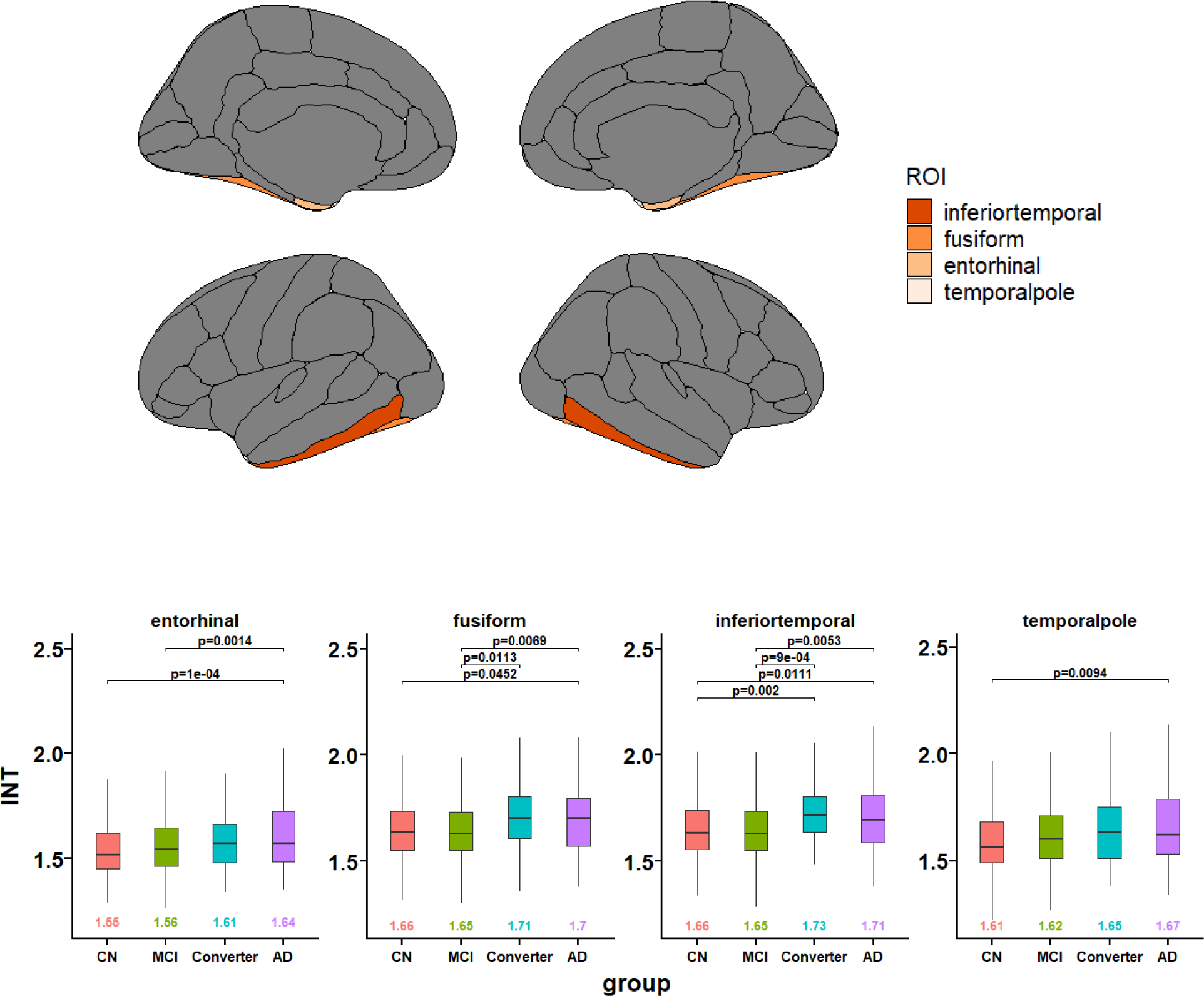
Cortical ROIs in which the INT values in AD group (and Converter group) are significantly longer than those in CN and MCI groups. Top: Brain visualization of the ROIs. The colormap indicate the significance of group differences in INT by ANOVA test (darker color indicates smaller p value). Bottom: The boxplots of the INT values by diagnosis groups on the identified ROIs. For each ROI, the mean INT values are shown by group on the bottom of the boxplot and the significant pairwise p-values are given on the top (threshold at 0.05). Age, sex, MFD and cortical thickness were controlled. Tukey method was applied to adjust for comparing a family of 4 estimates.

### 3.5 Significant cortical ROIs in which the MCI groups had shorter INT values than other groups

As shown in Figure 5, four ROIs had significantly lower INT values in the stable MCI group than the CN group, which are banks of the superior temporal sulcus (BanksSTS, CN-MCI: estimate= 0.051, Cohen’s *d* = 0.175, *t*(896) = 2.626, *p* = 0.0435), paracentral (CN-MCI: estimate= 0.050, Cohen’s *d* = 0.192, *t*(896) = 2.882, *p* = 0.0211), postcentral (CN-MCI: estimate= 0.048, Cohen’s *d* = 0.182, *t*(896) = 2.733, *p* = 0.0324) and precentral (CN-MCI: estimate= 0.036, Cohen’s *d* = 0.174, *t*(896) = 2.601, *p* = 0.0465) areas. Among them, significant differences between MCI and AD groups were found in the postcentral (MCI-AD: estimate= *−*0.070, Cohen’s *d* = *−*0.177, *t*(896) = *−*2.656, *p* = 0.0401) and precentral (MCI-AD: estimate= *−*0.069, Cohen’s *d* = *−*0.224, *t*(896) = *−*3.354, *p* = 0.0046) areas.

**Figure 5:**
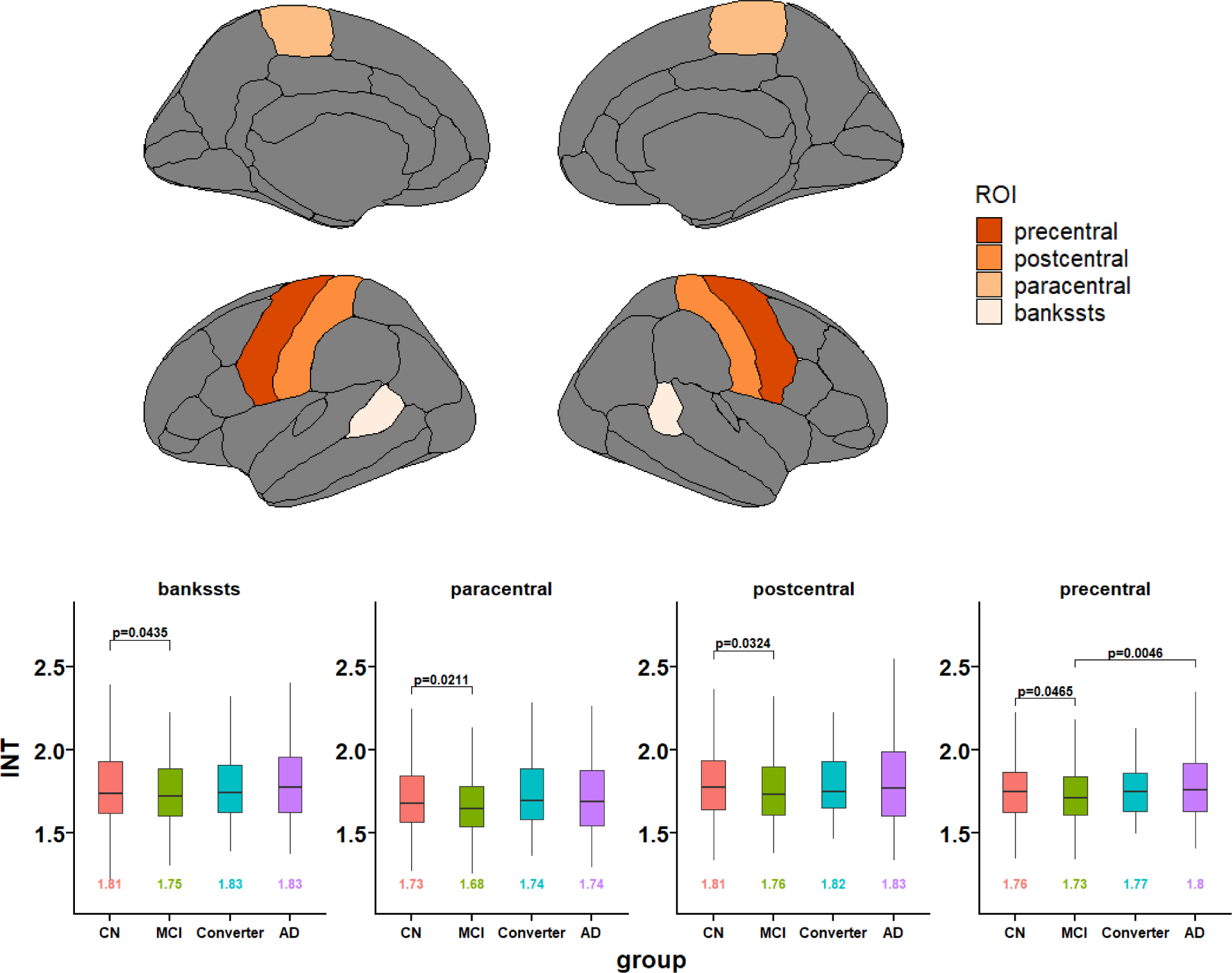
Cortical ROIs in which the INT values of MCI group are significantly shorter than CN group. Top: Brain visualization of the ROIs. The colormap indicate the significance of group differences in INT by ANOVA test (darker color indicates smaller p value). Bottom: The boxplots of the INT values by diagnosis groups on the identified ROIs. For each ROI, the mean INT values are shown by group on the bottom of the boxplot and the p-value of the significant pairwise difference is given on the top (threshold at 0.05). Age, sex, MFD and cortical thickness were controlled. Tukey method was applied to adjust for comparing a family of 4 estimates.

As shown in Figure 6, five ROIs were identified as having significantly shorter INT values in the MCI group than the AD group, which includes caudal middle frontal (MCI-AD: estimate= *−*0.068, Cohen’s *d* = *−*0.193, *t*(896) = *−*2.891, *p* = 0.0205), lingual (MCI-AD: estimate= *−*0.078, Cohen’s *d* = *−*0.219, *t*(896) = *−*3.283, *p* = 0.0059), middle temporal (MCI-AD: estimate= *−*0.065, Cohen’s *d* = *−*0.200, *t*(896) = *−*3.002, *p* = 0.0146), pericalcarine (MCI-AD: estimate= *−*0.100, Cohen’s *d* = *−*0.188, *t*(896) = *−*2.812, *p* = 0.0255), and superior temporal (MCI-AD: estimate= *−*0.068, Cohen’s *d* = *−*0.215, *t*(896) = *−*3.217, *p* = 0.0073) areas.

**Figure 6:**
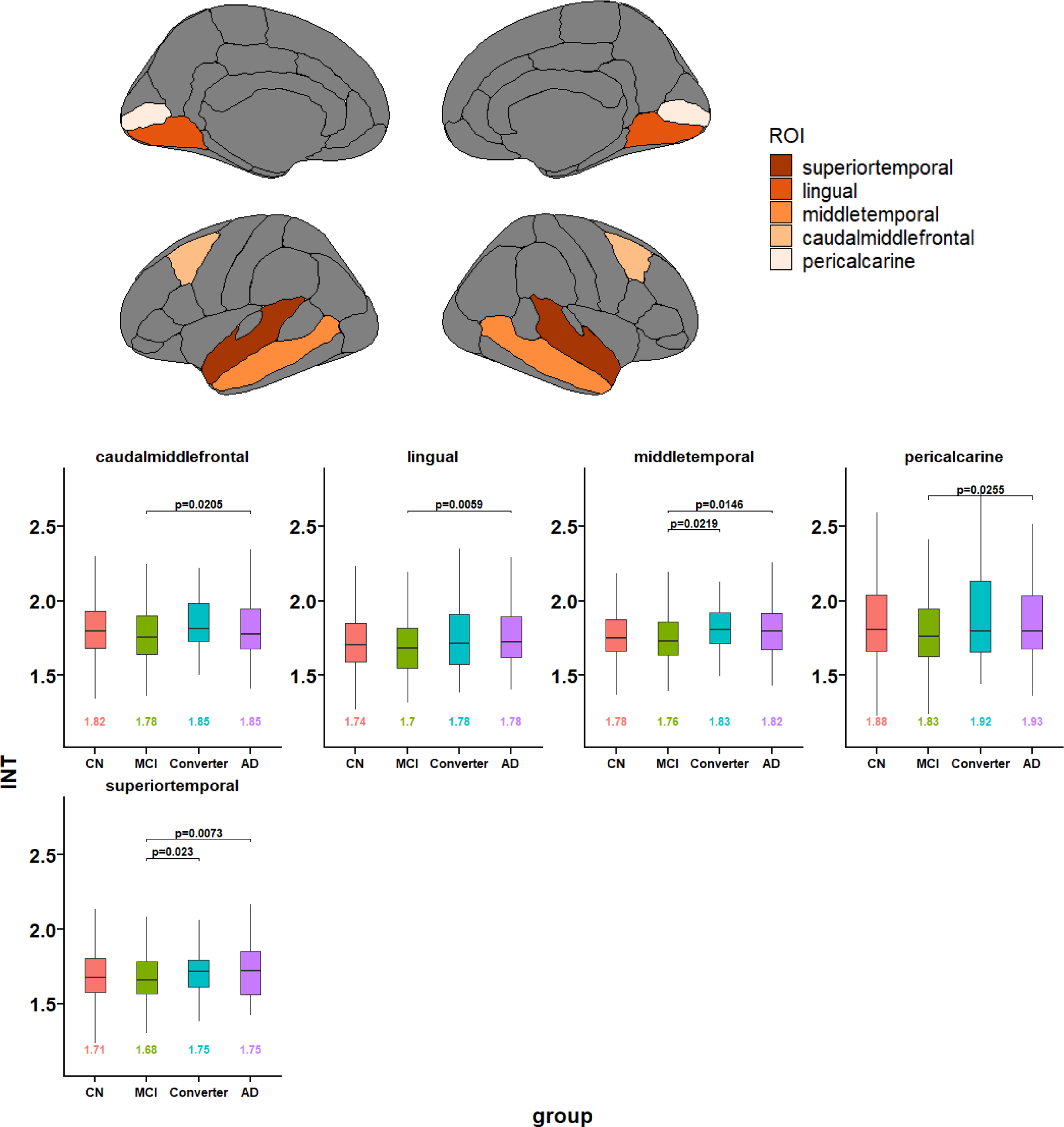
Cortical ROIs in which the INT values of MCI group are only significantly shorter than AD (and Converter). Top: Brain visualization of the ROIs. The colormap indicate the significance of group differences in INT by ANOVA test (darker color indicates smaller p value). Bottom: The boxplots of the INT values by diagnosis groups on the identified ROIs. For each ROI, the mean INT values are shown by group on the bottom of the boxplot and the p-value of the significant pairwise difference is given on the top. Age, sex, MFD and cortical thickness were controlled. Tukey method was applied to adjust for comparing a family of 4 estimates.

### 3.6 INT differences in subcortical regions

We investigated INT differences in 8 subcortical regions, including the hippocampus, a structure known to be compromised functionally and structurally in AD. As shown in Figure 7, four ROIs had significant group differences, which are caudate, hippocampus, pallium, and putamen. In general, the AD and Converter groups tended to have longer INT values than the CN and MCI groups. The contrasts are particularly apparent between the MCI and Converter groups (MCI-Converter, caudate: estimate= *−*0.057, Cohen’s *d* = *−*0.243, *t*(896) = *−*3.643, *p* = 0.0016, hippocampus: estimate= *−*0.042, Cohen’s *d* = *−*0.195, *t*(896) = *−*2.924, *p* = 0.0185, pallidum: estimate= *−*0.041, Cohen’s *d* = *−*0.210, *t*(896) = *−*3.147, *p* = 0.0092).

**Figure 7:**
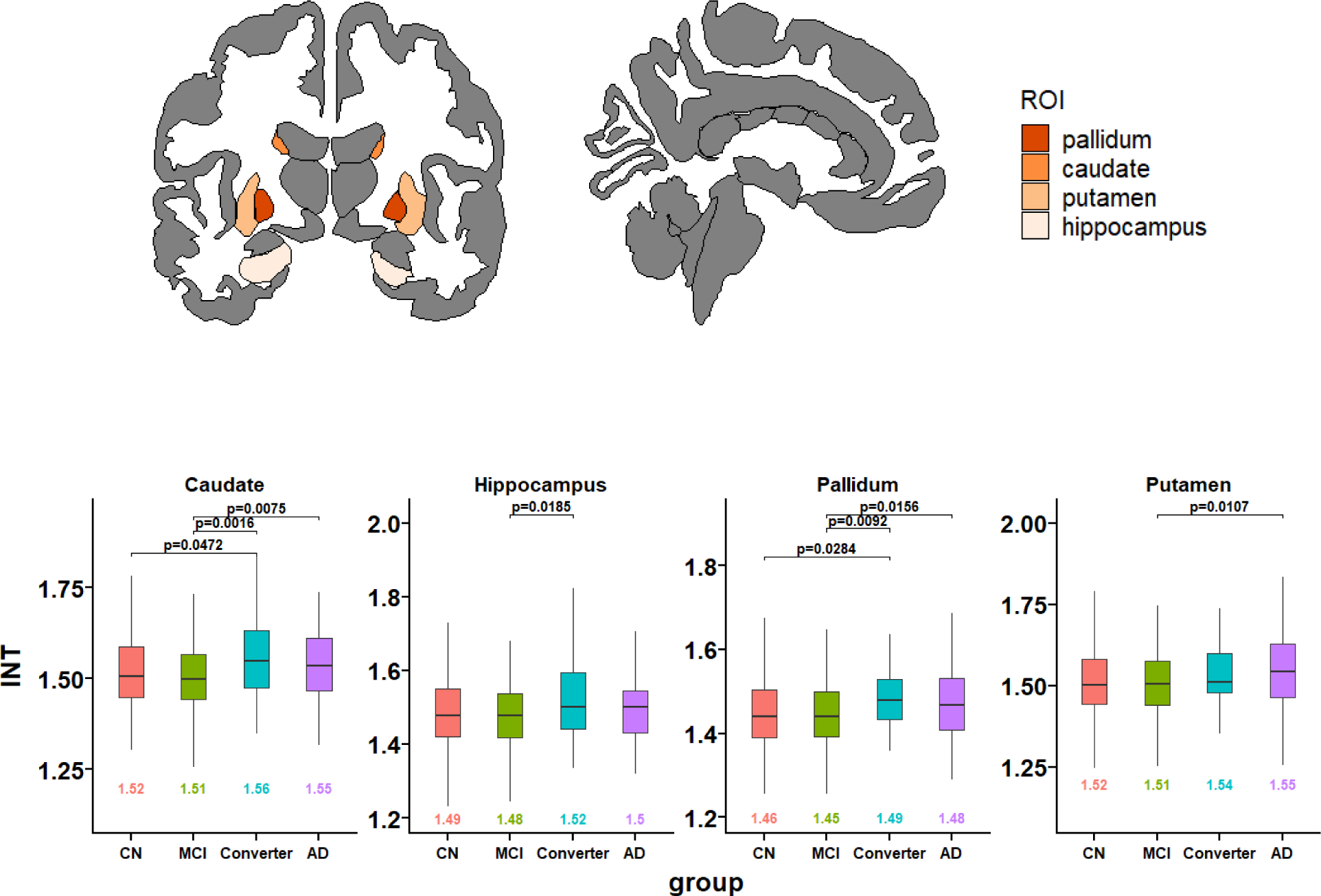
Subcortical ROIs in which the INT values have significant pairwise group differences. Top: Brain visualization of the ROIs. The colormap indicate the significance of group differences in INT by ANOVA test (darker color indicates smaller p value). Bottom: The boxplots of the INT values by diagnosis groups on the identified subcortical ROIs. For each ROI, the mean INT values are shown by group on the bottom of the boxplot and the p-value of the significant pairwise difference is given on the top. Age, sex and MFD were controlled. Tukey method was applied to adjust for comparing a family of 4 estimates.

### 3.7 Converter Group Shows Similar INT Profile as AD Group

In Figure 8, the INT contrast is minimal between AD and Converter groups, but mostly distinct between MCI and MCI groups. As shown in Figure 4, 5, 6 and 7, no significant differences have been found between AD and Converter groups. For the five ROIs that showed significantly longer INTs in the Converter group than the MCI group, four of them also had significantly longer INTs in the AD group (except for hippocampus). This includes fusiform (MCI-Converter: estimate= *−*0.068, Cohen’s *d* = *−*0.206, *t*(896) = *−*3.084, *p* = 0.013), inferior temporal (MCI-Converter: estimate= *−*0.081, Cohen’s *d* = *−*0.253, *t*(896) = *−*3.784, *p* = 0.0009), middle temporal (MCI-Converter: estimate= *−*0.077, Cohen’s *d* = *−*0.192, *t*(896) = *−*2.868, *p* = 0.0219) and superior temporal (MCI-Converter: estimate= *−*0.075, Cohen’s *d* = *−*0.190, *t*(896) = *−*2.852, *p* = 0.023) areas. Three ROIs have longer INT values in Converter group than CN groups, which are inferior temporal (CN-Converter, estimate= *−*0.074, Cohen’s *d* = *−*0.240, *t*(896) = *−*3.590, *p* = 0.002), caudate (CN-Converter, estimate= *−*0.040, Cohen’s *d* = *−*0.173, *t*(896) = *−*2.596, *p* = 0.047), pallidum areas (CN-Converter, estimate= *−*0.035, Cohen’s *d* = *−*0.186, *t*(896) = *−*2.778, *p* = 0.028).

**Figure 8:**
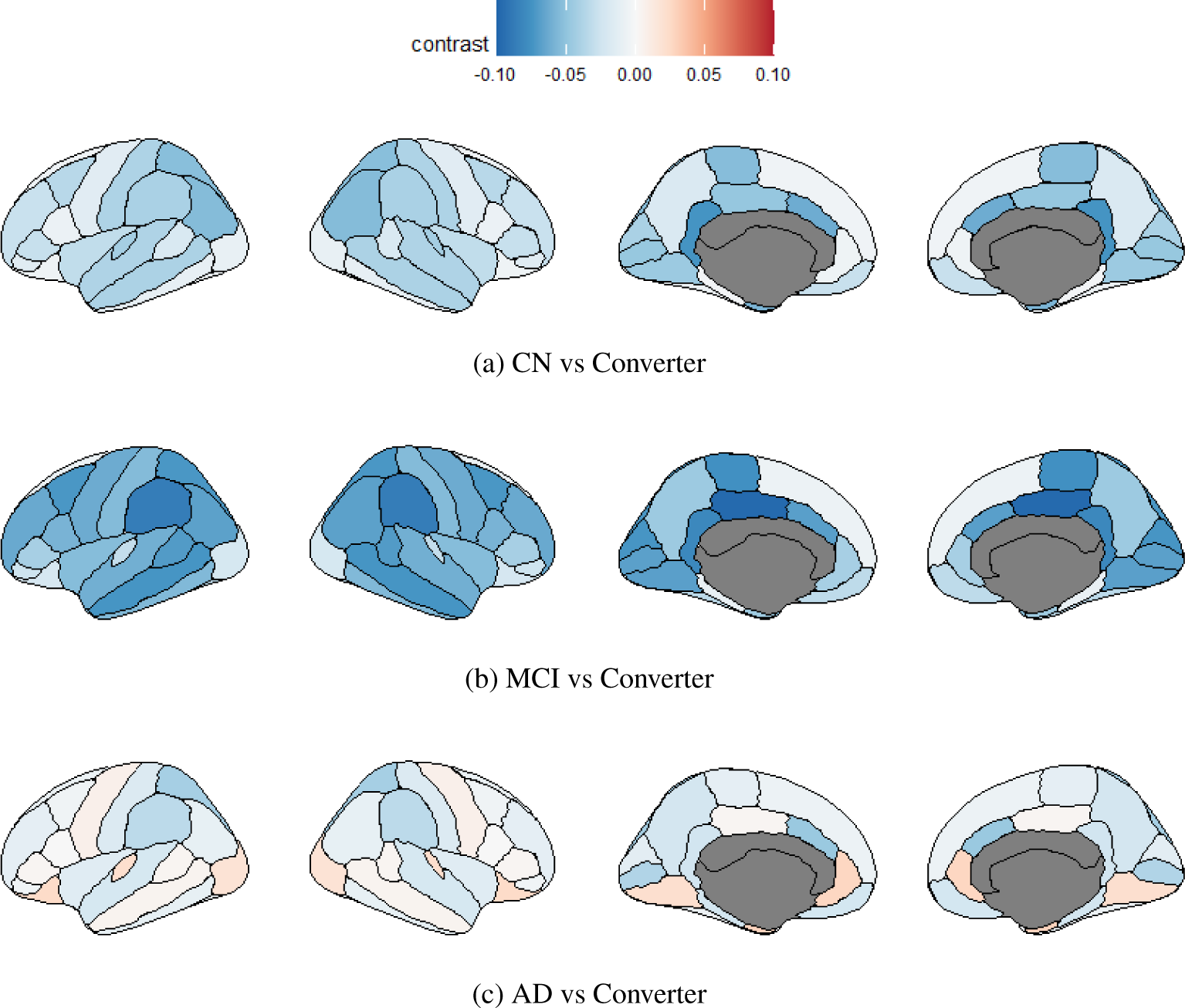
Contrast INT maps between Converter and CN, MCI, and AD groups, respectively. The effects of age, sex, MFD and cortical thickness were adjusted.

The pairwise comparison results and the effect sizes (mean INT values) indicate that the INT alterations in these areas in the Converter group are already very similar to the AD group at baseline, suggesting similar pathological changes prior to clinical presentation of AD.

## 4 Discussion

We used rs-fMRI data to investigate INTs in individuals of CN, MCI, including those who convert to AD, and AD populations. INT is closely related to brain functional hierarchy (13; 38). Using established cortical INTs from an independent young and healthy adult sample as a functional index of hierarchy, we found that 1) AD and Converter groups are similar, as they both had had longer INTs in low hierarchical order areas compared to CN; and 2) stable MCI is distinct from AD and Converter groups, which had a less pronounced hierarchical-gradient effect compared to CN with shorter INTs in high hierarchical order areas.

The pairwise comparison results showed that distinct cortical INT alterations in AD were present in the entorhinal, fusiform, inferior temporal, and temporal pole areas; subcortical alterations were in the caudate, hippocampus, pallidum and putamen. Three ROIs had significant INT alterations in the Converter group: the inferior temporal, caudate, and pallidum areas. Distinct cortical INT alterations in the MCI group were present in the banks of the superior temporal sulcus, paracentral, postcentral and precentral areas; specific differences between MCI and AD/Converter groups were present in the fusiform, inferior temporal, middle temporal and superior temporal areas.

Our results suggest distinct pathophysiological changes in the stable MCI and AD/Converter groups, at least as they relate to INT. For the AD and Converter groups, neural information is stored for a longer time in lower hierarchical order areas; while higher levels of the hierarchy seems to be preferentially impaired in stable MCI. Interestingly and perhaps counter-intuitively INT in primary somatosensory and motor cortices (BA 1, 2, 3, and 4, precentral and post central ROIs) and primary visual cortex (BA 17, pericalcarine) were lowest in the stable MCI group, perhaps an indication of non-AD aging changes in this group. The similarity in INT between the AD and Converter groups, and the differences between the MCI and Converter groups, suggests the potential for INT as a biomarker to predict conversion from MCI to AD. Future studies with larger samples of Converters are needed to develop INT — potentially in conjunction with other neuroimaging measures and/or cognitive and clinical panels — as a biomarker of conversion using multivariate machine learning methods.

Prior biophysical modeling work has indicated that INT depends on the strength of recurrent excitation within a brain region. Therefore, our observation of increased INT in the AD and Converter groups suggest increased E/I. Converging evidence suggests that E/I imbalance is a critical regulator of AD pathology (39; 40; 41). Studies shown that GABAergic dysfunction in AD impinges on the function of inhibitory neurons and on their ability to orchestrate and balance excitatory neurons (40), and an imbalance of excitation and inhibition will lead to epileptogenesis and AD pathogenesis (39; 22) — both of which point to increased E/I in AD. The similarity of INT values between the Converter and AD groups indicates that pathological E/I imbalance already started before the conversion to AD. The similarity of INT values between the Converter and AD groups indicated that pathological E/I imbalance predates diagnostic conversion to AD. Thus, identifying regions with disrupted E/I balance areas using INT from in AD and Converter groups constitutes an important first step to develop new diagnostic techniques and new treatment paradigms that aim to restore E/I imbalance for early intervention.

Altered INT was observed in several cortical areas implicated in the early stages of AD pathology. The entorhinal cortex is a medial temporal lobe brain region that often exhibits the earliest histological alterations in AD, including the formation of neurofibrillary tangles and cell death accompanied with deficits of memory and spatial navigation (42). The inferior temporal gyrus plays an important role in mediation of verbal fluency, such cognitive function is affected early in the onset of AD (43). Severe bilateral atrophy, neuropathologic changes and functional connectivity alterations have been found in the temporal pole areas of AD patients (44; 45; 46). Among subcortical regions, the hippocampus is the most extensively studied brain area in AD, with rapid neurodegeneration in the early stage of AD, which may be associated with the functional disconnection to other parts of the brain (47; 48). Interestingly, in healthy individuals the hippocampus has a low hierarchical order, indicating that its INT is short. This finding also demonstrates that despite substantial structural atrophy, INT is not shortened, but rather lengthened compared to controls. Abnormalities in INT in the basal ganglia were also identified. While underappreciated, studies have found atrophy of the caudate nucleus and other parts of the basal ganglia in the preclinical and clinical stages of AD (49; 50; 51). Consistent with current literature, these areas with altered INT may represent a stable biomarker for clinical diagnosis and an important therapeutic target in AD.

One strength of the study is that it is the largest rs-fMRI sample from ADNI in the literature. With a sample size of 945 it is well-powered to detect INT alterations both in terms of hierarchical gradient and individual regions. Furthermore, recent work has suggested that INT may have greater precision (i.e., test-retest reliability) as a measurement than other resting state network metrics, an important point in favor of its use, especially to the extent that it makes the work more replicable (15). With respect to limitations, the sample is largely Caucasian and may or may not generalize to other ethnic groups. Additionally, our study was intentionally limited to investigating the effects of global severity (CN and AD) and progression/stable MCI subtypes. Future work will focus on understanding the relationships between INT and other AD biomarkers, including *Aβ* and p-tau and its effect on cognitive test performance.

## Acknowledgements

Our work is partially funded by National Institute on Aging (R01AG062578), National Institute of Mental Health (R01MH129395, K01MH122774).

